# Wikipedia as a gateway to biomedical research: The relative distribution and use of citations in the English Wikipedia

**DOI:** 10.1101/165159

**Authors:** Lauren A. Maggio, John M. Willinsky, Ryan M. Steinberg, Daniel Mietchen, Joseph L. Wass, Ting Dong

**Affiliations:** Department of Medicine, Uniformed Services University of the Health Sciences. Bethesda, Maryland, United States of America.; Graduate School of Education, Stanford University, Stanford, California, United States of America.; Lane Medical Library, Stanford Medicine, Stanford California, United States of America.; Data Science Institute, University of Virginia, Charlottesville, Virginia, United States of America.; Crossref, Oxford, Oxfordshire, United Kingdom.

## Abstract

Wikipedia is a gateway to knowledge. However, the extent to which this gateway ends at Wikipedia or continues via supporting citations is unknown. Wikipedia’s gateway functionality has implications for information design and education, notably in medicine. This study aims to establish benchmarks for the relative distribution and referral (click) rate of citations—as indicated by presence of a Digital Object Identifier (DOI)—from Wikipedia, with a focus on medical citations.

DOIs referred from the English Wikipedia in August 2016 were obtained from Crossref.org. Next, based on a DOI’s presence on a WikiProject Medicine page, all DOIs in Wikipedia were categorized as medical (WP:MED) or non-medical (non-WP:MED). Using this categorization, referred DOIs were classified as WP:MED, non-WP:MED, or BOTH, meaning the DOI may have been referred from either category. Data were analyzed using descriptive and inferential statistics.

Out of 5.2 million Wikipedia pages, 4.42% (n=229,857) included at least one DOI. 68,870 were identified as WP:MED, with 22.14% (n=15,250) featuring one or more DOIs. WP:MED pages featured on average 8.88 DOI citations per page, whereas non-WP:MED pages had on average 4.28 DOI citations. For DOIs only on WP:MED pages, a DOI was referred every 2,283 pageviews and for non-WP-MED pages every 2,467 pageviews. DOIs from BOTH pages accounted for 12% (n=58,475) of referrals, making determining a referral rate for BOTH impossible.

While these results cannot provide evidence of greater citation referral from WP:MED than non-WP:MED, they do provide benchmarks to assess strategies for changing referral patterns. These changes might include editors adopting new methods for designing and presenting citations or the introduction of teaching strategies that address the value of consulting citations as a tool for extending learning.

## Introduction

Wikipedia, an online encyclopedia, has been described as a “gateway through which millions of people now seek access to knowledge” [1]. What has yet to be established is the extent to which the access to knowledge ends with Wikipedia. Increasingly, through the great “[citation needed]” movement of Wikipedia, this encyclopedia has become a potential gateway to the sources and more advanced forms of that knowledge. This paper addresses the question of the extent to which readers use those citations, as that bears on the well-documented turning to Wikipedia by healthcare providers and trainees [2-3]. There are obvious advantages of encouraging professionals to see Wikipedia as an effective gateway to learning from the literature as well as a ready reference for timely inquiry. Thus, this initial benchmarking study of readers’ referral rates for medical entries in Wikipedia is considered in relation to the work as a whole.

### Background

Launched in 2001, Wikipedia was envisioned as providing “every single person on the planet free access to the sum of all human knowledge” [4]. Some sixteen years later, the English version of Wikipedia has more than 250 million page views per day [5]. The number of references or citations on those pages has been increasing, with Wikipedia editors encouraged to include citations with links to reference sources to substantiate edits and enhance the overall verifiability of topic entries.

To support and encourage editors to include citations, Wikipedia provides guidelines for selecting references and templates for effectively adding citations to pages [6]. Footnote numbers are added to the text, typically at the end of a sentence making a claim, and lead to a bibliographic citation in the Notes section of the page. With medical pages, the citations are often to the biomedical research literature, and as such, the bibliographic information typically includes a hyperlinked Digital Object Identifier (DOI), PubMed ID, and less often a PubMed Central (PMC) ID. Each of these links connects readers to the research article abstract, with a further link to the full text. Full text is available either through subscription access, personal payment or open access, in the case of those articles with a PMC ID. PMC is a repository of open access articles that have been deposited by journals and authors, often in compliance with the NIH Public Access Policy [7]. Citations provide a valuable layer of verification for a Wikipedia page, if the page is treated as an end in the representation of knowledge on the topic, or the citations can be treated as gateways or entry points to a much vaster realm of research and inquiry, a growing portion of which – approaching half of the current literature – is publicly accessible [8].

Researchers have described multiple motivations for viewing Wikipedia content, such as a desire for information to fuel intrinsic learning, to meet work and/or school needs, and to make personal decisions [9]. While these motivations can often be satisfied by the topical overview provided by the typical Wikipedia entry, we propose that for these three specific information needs, Wikipedia’s function as a gateway to scholarly literature is potentially just as valuable a service. Furthermore, we contend that when it comes to health trainees’ and professionals’ use of Wikipedia, this gateway to the research literature function is of considerable consequence.

### A case for health information

Wikipedia has been identified as a major information resource for individuals seeking and engaging with health information [10]. In 2013, there were over 6.5 billion visits to medical content on Wikipedia [3]. Medical content on Wikipedia is overseen by WikiProject Medicine (WP:MED) [11]. WP:MED is under the purview of a highly active and well-coordinated volunteer project within Wikipedia and its sister sites that focuses on the coverage of medical topics. As a subset of Wikipedia, WP:MED pages account for approximately 1 percent of Wikipedia pages. In turn, healthcare professionals and trainees are known to be regular users of Wikipedia in clinical practice [12-13].

This use of Wikipedia bears on the practice of evidence-based medicine (EBM), in which physicians have been trained and encouraged over the last three decades to consult the “best available” research evidence in caring for their patients [14]. In the course of patient care, clinicians access and apply health information [15-16], including from Wikipedia [13, 17]. While Wikipedia entries can provide valuable overviews of medical topics, they do not, by design, provide the details of research studies needed to fully inform clinicians’ decisions about patient care [18]. For example, the English-Wikipedia writing guidelines prescribe against including drug dosages in medical entries and remind editors that Wikipedia is not a “how to manual” for practicing medicine [19].

This interest in evidence is where Wikipedia’s gateway function comes into play. For example, the pulmonary embolism entry in Wikipedia addresses therapeutic treatments broadly, but cites the Cochrane systematic review on the specific use of heparins and their dosage [20]. This Cochrane systematic review, which is considered an EBM “gold standard,” provides practicing clinicians who have subscription access to the Cochrane Library with the information necessary for effective treatment.

Similarly, for patients and their loved ones, Wikipedia entries provide overviews of medical topics, while also serving as a starting point for more advanced searches for health information tailored, for example, to their own characteristics (e.g., age, sex, race, and specific goals for their care) [21]. This information can play a role in shared health decisions with their physician, as has been shown for patients battling cancer, living with chronic illness, or deciding to undergo an invasive procedure [22].

This study intends to lay a foundation for analyzing and improving Wikipedia’s gateway-to-research function by determining the extent to which people currently use the citation links to the research literature (as marked by DOIs). Given that Wikipedia entries are consulted by a large class of health professionals, trained in research and encouraged to pursue EBM, and by motivated patients and their loved ones concerned with health issues, this study focuses on coverage of medical topics.

This initial study and those that follow are aimed at assisting in and improving

(a) the understanding of citation usage and function for different readers;

(b) the use of Wikipedia in professional education initiatives;

(c) the citational work of Wikimedia editors; and

(d) Wikipedia standards and systems for bibliographic entries to facilitate this gateway function.

## Methods

This is a descriptive study that aims to establish the rate of DOI referrals or clickthroughs from Wikipedia. Data for this study is specific to only pages with at least one DOI in the English version of Wikipedia for the month of August 2016. The aggregated data used for this analysis provides no indications of individual behaviors or identifiers, and thus did not require Institutional Review Board approval.

### Data Collection

This study utilizes the DOI assigned to the articles that appear in the research literature, as maintained by Crossref, as a persistent, resolvable, unique identifier to measure the presence and activities associated with the distribution and use of research citations in Wikipedia (Fig 1). When referred (or activated or clicked) by human or machine (web crawling spiders), the DOI link leads to a DOI link resolver. The DOI link resolver redirects to the publisher’s website, and specifically the article’s page with its abstract and metadata, with a link, typically, to the full text, or less often to the full text itself. A log entry of the click-through associated with the citation is recorded by the DOI link resolver.

**Figure 1.**
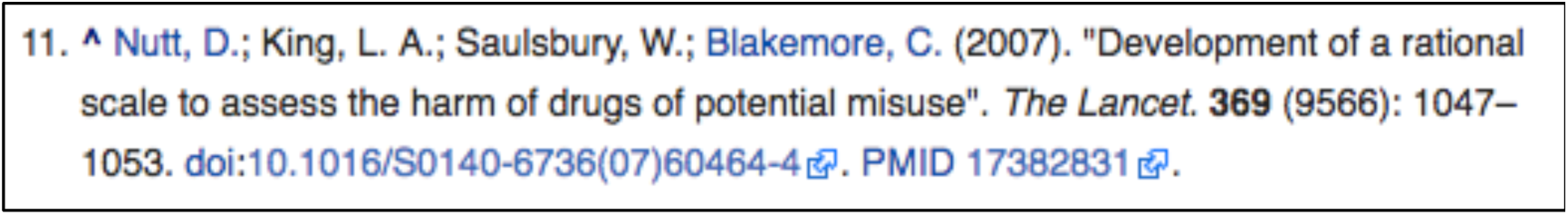
A citation with a DOI link from the References section of the Wikipedia page on Drug Prohibition Law to a 2007 scholarly article. The “doi” in the link acts as https://dx.doi.org when referred (clicked).

We accept that, like any other URL, bots may visit DOIs. The data set is filtered to include only resolutions where the agent sent a referral header, which excludes a large amount of automatic access. The research is predicated on two assumptions. Firstly, we assume that the presence of the referrer header indicates the presence of a human in the majority of cases. Secondly, we assume that the rate at which bots visit DOI citations is consistent across the categories of Wikipedia pages that we are comparing, which allows us to treat any bot activity as background noise and enables us to make relative comparisons between referrals from the categories.

Crossref, the largest DOI Registration Agency for scholarly content, provided access to August 2016 logs, which contain the DOI, date, time of resolution, IP address, and referrer URL. In October 2016, JLW processed the log entries, truncating the referring URLs to only include the full domain name (e.g., en.wikipedia.org) and the referring URL (including whether it was HTTPS or HTTP). Lastly, JLW filtered log entries to keep only resolutions with the domain name en.wikipedia.org and removed all sensitive information such as the IP address and precise time of referral to preserve user anonymity (Table 1; source code to process Crossref data: https://doi.org/10.5281/zenodo.822636 [23]).

**Table 1:** Example of data elements provided by Crossref for each DOI resolved in August 2016.

| Element | Example |
| --- | --- |
| Resolution Date | 2016-08-01 |
| DOI | 10.1002/14356007.a06_233.pub2 |
| Full referring domain name: | en.wikipedia.org |
| Subdomain of referring domain name: | en |
| The domain name: | wikipedia.org |

The Crossref data received did not include the name of the Wikipedia page from which the referral originated. Therefore, the Crossref data alone was insufficient to determine whether the referral originated from a link on an entry covering a medical topic. To ascertain the possible origins of the referral, for each Wikipedia page that contained at least one DOI, we collected the following: page ID; namespace (Wiki page type); page title; external link to doi.org; and whether it was a WP:MED page (Table 1). We defined a page as WP:MED if it was contained in at least one of the 22 WikiProject Medicine categories [24] and included associated Talk pages. Data was collected for each day in August 2016. Additionally, DOIs were classified as coming from one of three locations:

1) WP:MED --The DOI citation appears only on a medical topic page.

2) Non-WP:MED --The DOI citation does not appear on a medical topic page.

3) BOTH --The DOI citation appears on both WP:MED and non-WP:MED pages.

(Source code to extract, summarize and categorize Wikipedia data available here: https://doi.org/10.5281/zenodo.824813)[25].

In February 2017, we merged the Crossref data (DOIs referred in August) with the data extracted from Wikipedia pages for each day of August. The merge revealed from which Wikipedia page(s) the DOI may have been referred. For example, on August 3, the DOI (http://dx.doi.org/10.1016/S0140-6736(07)60464-4) associated with the *Lancet* article entitled “Development of a Rational Scale to Assess the Harm of Drugs of Potential Misuse” [26](see fig. 1), which was referred three times, was on that date on a WP:MED page (“Substance Abuse”) and a non-WP:MED page (“Drug Prohibition Law”) and thus classified as BOTH.

As Wikipedia is a dynamic resource, we also examined the change in number of Wikipedia pages with a DOI and the total number of DOI citations present over the study period. In April 2017, we created a summary file of DOI references and pages present for each day in August 2016. Each DOI reference and Wikipedia page were classified as WP:MED, non-WP:MED or BOTH and summed to determine the relative distribution of DOI references and Wikipedia pages present.

Using the Wikipedia Pageview API (application programming interface) [27], we extracted pageview data for Wikipedia pages with at least one DOI citation including all WP:MED pages with at least one DOI citation. This included pageviews identified as coming from either desktop or mobile devices, while excluding most traffic from spiders, which consisted of, during our study period, about 14 percent of overall English Wikipedia pageviews, with approximately 6 percent of visits to pages with DOI references, 4 percent of visits to WP:MED pages with DOIs.

The data was primarily analyzed using descriptive statistics. To examine whether there is any difference between the referral rates of WP:MED and non-WP:MED, we performed independent sample t-tests.

## Results

The results of this analysis are for English Wikipedia pages with at least one DOI through the month of August 2016. Over the course of the month, the daily average of Wikipedia pages was 5.2 million, of which 4.42 percent (n=229,857) included at least one DOI citation [28]. 68,870 pages were under the purview of WP:MED, with 22.14 percent (n=15,250) of them featuring one or more DOI citations. WP:MED has more pages with DOI citations per page than the rest of Wikipedia. These WP:MED pages also contain twice as many DOI citations, with an average of 8.88 citations per page, compared to those outside of WP:MED, with 4.27 DOI citations per page (Table 3). This establishes what might be thought of as the higher saturation level in the citation of scientific research articles among WP:MED pages compared to the whole, while also attesting, more generally, to how pages with a DOI citation tend to include clusters of them.

**Table 2.** Data elements extracted for each English-language Wikipedia page that included at least one DOI

| Elements | Example |
| --- | --- |
| Page ID | 207165 |
| Namespace | Main |
| Is WikiProject Medicine | True |
| Page title | Pulmonary embolism |
| Link to doi.org | https://dx.doi.org/10.1002/14651858.CD001100.pub3 |

**Table 3.** Average number of DOI citations and average number of pages with DOIs for WP:MED and non-WP:MED for August 2016.

|  | WP:MED | Non-WP:MED | TOTAL |
| --- | --- | --- | --- |
| DOI citations | 134,459 (13%) | 910,624 (87%) | 1,043,614 (100%) |
| Pages w/ DOIs | 15,143 (7%) | 212,746 (93%) | 228,178 (100%) |
| DOI citations/page | 8.88 | 4.28 | 4.57 |

In terms of change over the course of August, the number of pages with at least one DOI citation increased at a significantly higher rate for Non-WP:MED than for WP:MED [*t* (58) = 2.158, *p*=.035, with a medium effect size (Cohen’s *d* =.557)] (Table 4). Although fewer pages with DOI citations were being added to WP:MED in August, this section of Wikipedia kept up in the number of DOI citations added. WP:MED added 1,872 DOI citations for an increase of 1.42 percent in August, while non-WP:MED added a comparable number with 13,066 (1.44%) DOI citations [t (58) = -.265, p=.792] (Table 5).

**Table 4.** Increases in total Wikipedia pages with one or more DOI citations for WP:MED and non-WP:MED during August 2016.

|  | WP:MED | Non-WP:MED | Total Pages |
| --- | --- | --- | --- |
| August 31, 2016 | 15,250 (7%) | 214,607 (93%) | 229,857 (100%) |
| August 1, 2016 | 15,064 (7%) | 210,961 (93%) | 226,025 (100%) |
| August page gain<sup>a</sup> | 186 (1.2%) | 3,646 (1.7%) | 3,832 (1.6%) |

<sup>a</sup> Percentage gain is from August 1, 2016 start date.

**Table 5:** Number of DOI citations that appeared in WP:MED and non-WP:MED during August 2016.

|  | WP:MED | Non-WP:MED | TOTAL |
| --- | --- | --- | --- |
| August 31, 2016 | 134,181 (13%) | 914,844 (87%) | 1,049,025 (100%) |
| August 1, 2016 | 132,309 (13%) | 901,838 (87%) | 1,034,147 (100%) |
| August DOI gain | 1,872 (1.42%) | 13,066 (1.44%) | 14,878 (1.44%) |

As for the referrals or use readers made of DOI citations, for those DOI citations that only appeared on WP:MED pages, a reader made a referral for every 2,283 pages that users viewed. For those DOI citations that only appeared on non-WP-MED pages, readers made a referral every 2,467 pages viewed during August 2016. These results do not include the 58,475 referrals for those DOI citations that could be found in BOTH. Not being able to identify the source page for 12 percent of the referrals limits the confidence that can be placed in the similarity of referral rates between WP:MED and non-WP:MED. While this result can still be used as a benchmark for later assessing relative changes in the referral rate within WP:MED, it does not currently offer evidence as to whether readers are referring the citations with DOIs any more frequently in WP:MED than elsewhere.

A substantial portion of DOIs were referred at least once, with the average citation receiving multiple referrals (Table 7). For example, 32 percent of DOI citations exclusive to WP:MED were referred during August, while on non-WP:MED pages, 24 percent of DOI citations exclusive to those pages were referred. The DOI citations appearing in BOTH, not surprisingly, had a higher proportion of referrals (52%). For those DOI citations that were referred, the average frequency of referral for WP:MED pages was 2.19 times, for non-WP:MED pages 2.56 times, and again a higher rate of 4.23 times for those citations that appeared on BOTH pages (Table 8). Again, these results offer a potential benchmark for assessing relative changes in referrals within WP:MED, but no evidence on whether WP:MED is an area of greater or even possibly less referral activity than the rest of Wikipedia.

**Table 6.** Pageviews of pages with at least one DOI citation and the referrals from DOI citations during August 2016.

|  | WP:MED | Non-WP:MED | BOTH | Total |
| --- | --- | --- | --- | --- |
| Pageviews w/ DOI | 145,246,622<br>(14%) | 875,180,267<br>(86%) | - | 1,020,426,889<br>(100%) |
| DOI referrals made | 60,958 (13%) | 354,706 (75%) | 58,475<br>(12%) | 474,140 (100%) |
| Pageviews/referral | 2,283 | 2,467 | - | 2,152 |

**Table 7.** Unique DOI citation referrals as a proportion of unique DOI citations for WP:MED alone, in non-WP:MED alone, and for BOTH during August 2016.

|  | WP:MED | Non-WP:MED | BOTH | Total |
| --- | --- | --- | --- | --- |
| Unique DOI citations | 87,932 (13%) | 567,482 (83%) | 26,787 (4%) | 682,203 (100%) |
| Unique DOI referrals | 27,840 (15%) | 138,393 (77%) | 13,826 (8%) | 180,060 (100%) |
| Percent DOIs referred | 32% | 24% | 52% | 26% |

**Table 8.**
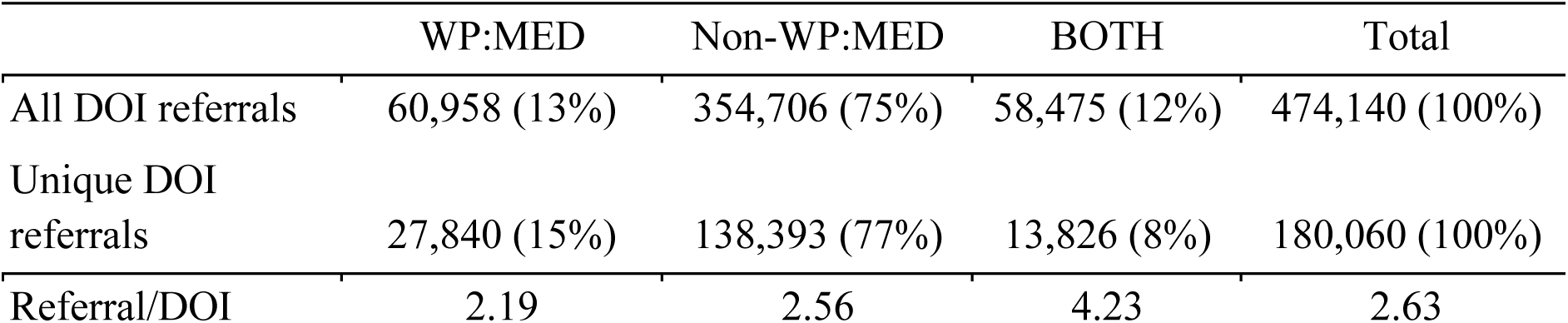
Average DOI citations, total referrals from DOI citations, and unique DOI citation referrals during August 2016.

This study also identified the most frequently referred DOI citations for August, 2016, providing a sense of what papers readers favored in pursuing Wikipedia’s gateway effect. The list was led by the 1999 *Science* article “Association of BRCA1 with the hRad50-hMre11-p95 complex and the DNA damage response,” which had 6,653 referrals [29]. It was cited in BOTH (i.e., WP:MED and/or non-WP:MED pages). This popular citation points to a further caution in such studies. The article is cited in four pages, three related to low-traffic pages with technical descriptions of proteins generated by the ProteinBoxBot, which is a program for the automated creation of such Wikipages. For August, these three pages had a total of 1,035 pageviews [27]. The fourth Wikipedia entry with this citation is “BRCA1,” which was viewed 15,628 times [27]. This suggests the majority of DOI referrals were generated from the BRCA1 page, which is a WP:MED page. Notably, this page had 1,090 pageviews on August 4. On this day, the news magazine, *New Scientist,* published the article, “Counting genetic mutations predicts how soon you’ll get cancer” [30], which prominently featured BRCA1 in its lead paragraph. This suggests that medical topics mentioned in news media may translate to Wikipedia pageviews and subsequent DOI referrals.

The second most referred citation, with 1,046 clicks, was “Volcanic Aerosols: The significance of volcanic eruption strength and frequency for climate” from a 2004 issue of *Quarterly Journal of the Royal Meteorological Society* [31]. It is cited on a non-WP:MED page. The third most frequently referred citation was to the 2014 article “Eye lens radiocarbon reveals centuries of longevity in the Greenland shark (Somniosus microcephalus)” in *Science* [32]. It was referred 354 times from 13 different Wikipedia entries, including the WP:MED entry on “Aging,” as well as one for the year 1634, which is has been suggested to possibly have been the birth year of “the Female Greenland shark (still alive in 21st century),” all of which speaks to the rich use of DOI citations.

In the case of DOI citations used in WP:MED alone, the most referred instance, ranked 31st with 155 referrals, was “Growth and ovarian function in girls with 48, XXXX karyotype-patient report and review of the literature,” published in the *Journal of Pediatric Endocrinology and Metabolism* [33]. This article is cited in the Wikipedia entry for the XXXX syndrome, and is listed under “Further reading.” The abstract for this 2002 article is freely available from PubMed, while access to the article itself costs $42.00 from its publisher De Gruyter, raising another set of issues to consider with Wikipedia’s gateway function. The frequency counts for individual DOI citations offers yet another benchmark for assessing the impact of different strategies on relative rates of referral of DOI citations in WP:MED.

## Discussion

The results of this study demonstrate what can be established about the distribution and use of scientific research citations, which have been assigned a DOI. This provides a helpful starting point in assessing the extent to which Wikipedia serves its users in an area such as WP:MED as a gateway to the sources of knowledge, whether for purposes of verification and/or further learning.. The methods utilized in this study include both a daily crawling during the month of August 2016 of Wikipedia to determine how many of its pages have citations with DOIs and how many such citations exist both inside and outside of WP:MED. A second method – involving the DOI resolution provided by Crossref – enabled a determination of which and how many of the DOI citations inside of or beyond WP:MED were referred (or clicked).

The results of merging and analyzing these two data sources indicate that the WP:MED pages are not characterized by greater gains over the month in pages or the number of citations added. They appear to have more DOI citations on them than pages outside of WP:MED that have at least one DOI citation. As to how often in a month these citations are referred, the data could not provide a definitive comparison of within and outside of WP:MED, because a good number of citations are to be found on both types of pages. Without being able to tell from which location in Wikipedia the BOTH referrals were made, no comparison can be drawn between the referrals from within WP:MED and from outside of it. At best, it can be said that these results did not provide evidence indicating that citations on WP:MED are more frequently explored (as they might have if the WP:MED were larger than non-WP:MED and the BOTH referrals combined). Given the evidence that health professionals use WP:MED and this community of users has reason to use Wikipedia as a gateway more frequently than other users, we find that there are good reasons for further inquiries.

To that end, the results of the present study provide a series of benchmarks that can be used to assess whether there are strategies for changing referral patterns. These changes might result from encouraging editors to adopt better methods for designing and presenting the citations. For example, a system of access icons was recently rolled out to help Wikipedia editors visually indicate if the full text of a citation is publicly accessible [34]. This indication of accessibility may increase referrals by encouraging users, such as physicians, that have admitted reluctance to click citation links due to the expectation of facing a paywall [35]. A second area to explore, initially on a smaller scale, is to introduce teaching strategies in graduate and continuing medical and health education that address the value of consulting the citations as a tool for extending learning. A third consideration is that changes may take place over time in user expectations and interests in accessing the sources to verify authorities and/or learn more about a topic (in an age of misinformation).

In the first instance, the Wikipedia guideline “Wikipedia:Citing sources” [36] provides extensive direction and tools for citing sources. But there is clearly room for improving the extent to which citations do more than signal verification by providing a channel to further learning and greater understanding. This may involve exploring the differences between citations in footnotes and “further reading” lists. It might include the multiple links associated with the citations in Wikipedia. For example, the typical citation considered here offers a hyperlinked article title (with an arrow icon), PMID (PubMed ID), DOI (with an arrow icon) and occasionally a PDF icon, arXiv ID or a Bibcode, as a further compact identifier. A subset of the citations in the life sciences also have a PMC link, along with a green open access icon, leading to PMC where the article is made freely available to readers. The citations include on rare occasions short annotations, with additional links, such as “lay summary.” While having three or four links per citation may seem to increase the likelihood of referral to the source, it may also deter use. Lastly, the variety of links and icons present highlights a study limitation in that we were only able to utilize DOI referral data, therefore missing the clicks to other resources, notably for medicine Pubmed and PMC. Ideally, future research will be able to include referral data from these resources.

## Conclusion

This study provides a number of measures for both the citing and use of research articles, as indicated by DOIs, in support of Wikipedia entries both inside WP:MED and outside of it. It also demonstrates a number of current limits in what can be ascertained about the use of the citations by readers (allowing for some referrals by bots). These limits arise from the need to protect the privacy of Wikipedia users. However, even with the use of completely anonymized data, there is still the need to establish the value of this benchmarking research in terms of contributing to the Wikipedia project. The present study has established that although there is a greater density of citations with DOIs on such pages in WP:MD, there is no evidence of any greater use or referral of those citations compared to those used outside of WP:MED. This study’s measures do afford a certain number of benchmarks on DOI citation usage for assessing the impact of strategies intended to increase referral within WP:MED, as well as changes in general behaviors in the use of Wikipedia.

Next steps for this program of research include refining the measurement techniques and strategies in ascertaining which citations are being referred within WP:MED, which will need to take place within Wikipedia and Wikimedia standards for such investigations. The research question remains whether there is greater educational value to be had in Wikipedia through the use of these citations, especially among health professionals. Or in user terms, to what extent do readers have additional capacity and interest in following through on the sources behind the citations if that process could be facilitated through improvements in information design; increases in open access to sources; and targeted educational efforts for health professionals?

## Acknowledgements

The authors would like to acknowledge Jake Orlowitz, the Head of the Wikipedia Library, for his early contributions in the conceptualization of this study and encouragement throughout the entire project.

## Data Deposit

Wass JL, Steinberg RM, Maggio LA. (2017). Data for Wikipedia as a gateway to biomedical research [Data set]. Zenodo. http://doi.org/10.5281/zenodo.831459

